# Neural and behavioral catalysts of ongoing memory retrieval

**DOI:** 10.1101/2025.03.28.646007

**Authors:** Matthew R. Dougherty, Anuya Patil, Katherine D. Duncan

**Affiliations:** Department of Psychology, University of Toronto, 100 St. George Street, 4th floor, Toronto, ON, Canada, M5S 3G3

**Keywords:** episodic memory, human, fMRI, retrieval, memory states, SN/VTA, basal forebrain, neuromodulators, reinstatement

## Abstract

Memory retrieval is notoriously variable. Various neurocognitive states have been theorized to affect retrieval success from moment to moment, but the presence and catalysts of these states in the human brain remain largely understudied. Building on previous work, we studied how recent memory judgments during an ongoing retrieval task (i.e. retrieval judgments vs. novelty detection) and their corresponding neural activity prepare the brain to reinstate unrelated memories during upcoming trials. High-level ventral stream regions, including the hippocampus, reinstated memories with significantly greater fidelity following recent retrieval judgments compared to novelty detection. Activity in dopaminergic nuclei rose during retrieval judgments, which predicted and partially mediated the effect of retrieval judgments on upcoming reinstatement. While dopaminergic nuclei activity predicted upcoming reinstatement, it did not predict upcoming retrieval accuracy. Exploratory analyses revealed the opposite effect in the dorsal and lateral prefrontal cortex, whose activity predicted upcoming retrieval accuracy, but not reinstatement. These results point to distinct neural contributions to what is reinstated and how it may guide memory decisions, and by identifying dopaminergic nuclei as partial mediators of reinstatement, our results open new avenues for investigating how neuromodulatory states may dynamically shape memory accessibility.

**Significance Statement:** Why can we effortlessly recall memories in some moments but struggle at others? Here, we draw on computational models to uncover why human brains are sometimes better prepared to remember and how to nudge them into that state. We discover that recent retrieval judgments, compared to recent novelty detection, increase upcoming accuracy and neural reinstatement of unrelated memories. This effect is so powerful that key memory regions only show reinstatement following preceding retrieval judgments. We found that dopaminergic nuclei are more active during retrieval judgments and predict upcoming memory reinstatement. This pattern partially explains why engaging in remembering prepares your brain to reinstate other memories and reveals new insights for the role dopaminergic nuclei may play in retrieval.

## Introduction

Memory retrieval is notoriously inconsistent. At times, past events vividly spring to mind with little effort; at other times, even well-learned associations prove stubbornly inaccessible. While the quality of initial memory formation contributes to this variability (1–5), long-standing theories (6) and neurocomputational models (7–10) propose that neurocognitive states established just before a retrieval attempt may also play a role. A key prediction from these frameworks is that ongoing acts of retrieval establish processing biases that facilitate access to other, even unrelated, memories. Behavioral studies have consistently supported this prediction (11–13), with recent work (13) showing that making a retrieval judgment on one trial improves associative retrieval accuracy on the next. However, whether these behavioral biases correspond to measurable changes in the neural mechanisms of memory, and what drives them, remains untested.

Extending the study of these biases to the brain is crucial for understanding whether they reflect changes in how, where, and what information the brain reinstates during acts of retrieval. Neocortical reinstatement of patterns during retrieval that were originally present during encoding, reliably indexed through Encoding-Retrieval Similarity (ERS) analyses (14–18), critically depends on the hippocampus (15, 19). The hippocampus operates in processing modes that switch between encoding novel information (pattern separation) and retrieving stored associations (pattern completion) (20–23). Because the mechanisms underlying the switch take time to unfold, states are theorized to temporally extend, biasing incoming information toward processing consistent with the already-established state (7–10). On this account, the same events that have been previously shown to bias behavioral markers of memory (11–13) should also bias hippocampal processing: retrieving one memory should establish a hippocampal pattern completion state, facilitating retrieval of subsequent unrelated memories, whereas detecting novelty would, comparatively, bias processing away from retrieval. As cortical ERS depends on the hippocampus (15, 19), it also follows that ERS should be relatively higher following retrieval of an unrelated memory compared to following novelty detection.

While ERS can illuminate whether these biases extend to the reinstatement of memories, it does not reveal their source. Neural activity during preceding memory judgments may work to establish these biases, particularly by driving the hippocampus into one processing state rather than another. Neuromodulators including acetylcholine (ACh), dopamine (DA), and norepinephrine (NE) are potential catalyst candidates. Each are released in response to memory-related functions including novelty detection (24–29), motivation (30–34), and attention (34–39), and they exert lingering behavioral effects on memory (33, 36, 39–43) and modulatory effects throughout the brain, including in the hippocampus (44). For example, rodent models propose that hippocampal ACh concentration biases the pattern completion/separation trade-off, with lower levels promoting pattern completion (43). Although ACh models provide the clearest mechanistic precedent, dopaminergic and noradrenergic systems are recruited by the same motivational and novelty factors, leaving open the possibility that other neuromodulators may establish the upcoming bias. While direct measurement of these neuromodulators is not possible using functional Magnetic Resonance Imaging (fMRI), we investigate the different candidates through univariate activity in the distinct nuclei that release them as a proxy of their engagement (45, 46).

Frontoparietal structures associated with initiation of the retrieval mode may also play a role in establishing lingering memory biases. Retrieval mode theories propose that episodic retrieval depends on a sustained goal-directed attentional orientation maintained by frontoparietal networks — one that directs processing toward internally stored representations and supports memory search (6, 47). This orientation is distinct from the reactivation of a stored trace itself, but may be necessary for the reactivated information to result in the full experience of a memory. As this orientation has been theorized to be sustained across time (48), strong activation in regions associated with the retrieval mode during one memory judgment may establish a state that benefits upcoming retrieval. While we did not design our task to isolate retrieval mode regions, we explored whether regions identified in other work (49–56) contributed to ongoing retrieval biases.

Here, we examined how different preceding memory judgments – reflecting retrieval or novelty detection – influenced upcoming recall and neural reinstatement during an ongoing associative memory retrieval task. Participants studied a set of associations, then at retrieval were shown either previously studied cues (prompting retrieval of the studied associate) or novel items (prompting novelty detection). We hypothesized that memory reinstatement (ERS) would be greater in the hippocampus and highly connected regions following preceding retrieval judgments compared to preceding novelty detection. We explored whether neuromodulatory ROI activity initiated this bias, with cholinergic basal forebrain activity as our primary *a priori* candidate. More exploratorily, we examined whether frontoparietal regions associated with recruiting and sustaining retrieval orientation contributed to the observed biases. Our results show that memory reinstatement is robustly modulated by preceding memory states, point to the contributions of dopaminergic nuclei in establishing this bias, and illuminate distinct neural contributions to behavioral and neural retrieval biases.

## Results

### Creating an fMRI paradigm suited to investigate how memory states affect upcoming reinstatement

We designed a task to evaluate how recent memory judgments, and the brain states they evoke, influence the upcoming neural reinstatement of memories. To balance temporal constraints associated with short-lived biases (9, 57) and blood-oxygen-level-dependent (BOLD) signals (58), we organized trials at retrieval into pairs and tested representationally distinct stimulus categories across trials, which allowed us to limit inter-trial intervals (ITIs) between paired trials to 1 second (Fig. 1a). Each pair’s first trial was treated as an induction of a neural state associated with either the detection of a trial-unique new object or the retrieval of a previously studied old object (*induction trial*). Each pair’s second (*probe*) trial tested how these states influenced successive memory retrieval accuracy and neural reinstatement through the presentation of new and old words, previously studied with scenes or faces. Participants made recognition judgments on both trials and, when recognizing items, attempted to recall associated information from encoding (see *SI Appendix: Materials & Methods* for full design specifications). Induction trials were classified according to subjective judgments: “new” responses indexed novelty detection, whereas any “old” responses indexed retrieval, regardless of associative success (see *SI Appendix: Materials & Methods* for response options). We extended ITIs between trial pairs to 5 seconds to increase reinstatement analysis sensitivity.

**Fig. 1.**
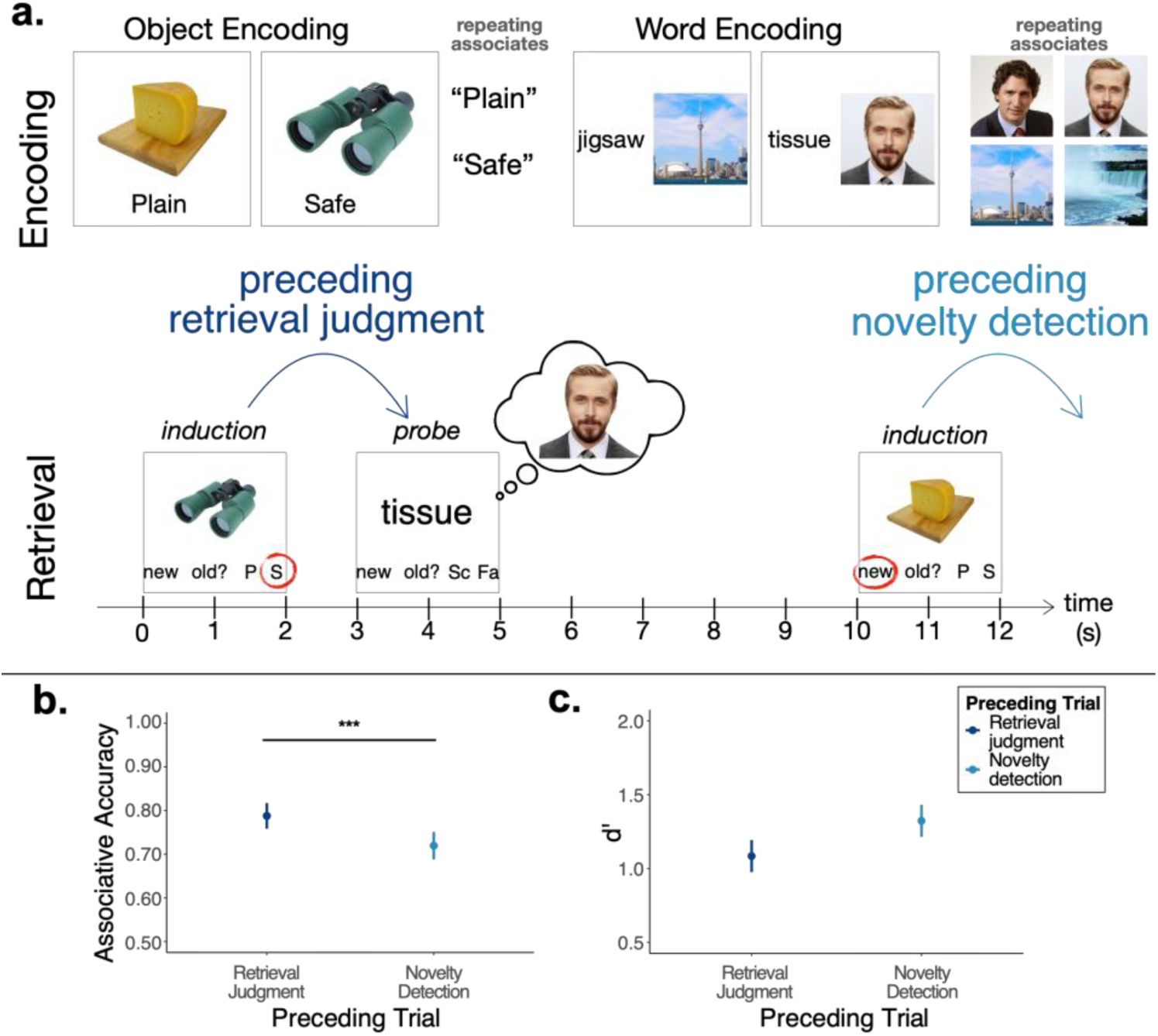
Task Design & Behavioral Analyses. **a**, Schematic of encoding & retrieval tasks. During Object Encoding, participants associated trial-unique objects with repeating words. During Word Encoding, participants associated trial-unique words with repeating face and scene images. At retrieval, participants were shown the trial-unique half of an encoded pair (object image or non-repeating word) and asked to retrieve its associate. Trials were structured in pairs, with object trials acting as memory state inductions and word trials acting as probes to measure neural reinstatement & associative memory accuracy (see *SI Appendix: Materials & Methods*). **b**, Associative memory accuracy on probe trials as a function of preceding memory experiences. Associative memory accuracy was calculated as the number of correct associative hits out of all associative attempts (i.e. responding with “Scene” or “Face”). Points represent the estimated marginal means, and error bars represent their standard error from a generalized linear mixed effects model (see *SI Appendix: Materials & Methods*). **c**, Item-only recognition memory (d’) on probe trials as a function of preceding memory experiences. Item-only recognition memory was calculated for all trials where participants did not respond with the correct associate, evaluating their ability to recognize previously seen words when associative recall was not successful. Points represent mean d’ across participants for trials preceded by retrieval judgments and novelty detection, respectively. Error bars represent the standard error of the mean, applying the Morey, 2008 correction for within-participant comparisons. *** indicates p < 0.001.

First, we assessed how our induction trial manipulation influenced memory retrieval accuracy. After replicating Patil and Duncan (2018), we assessed how induction trial judgments influenced successive neural reinstatement of memories using ERS analyses. We then measured how neuromodulatory nuclei responded on induction trials, before determining if these responses mediated the impact of recent memory judgments on neural reinstatement. Lastly, we explored whether frontoparietal regions associated with the retrieval mode played any role in establishing behavioral and neural biases in memory retrieval.

### Preceding memory judgments selectively influence associative retrieval accuracy

We first sought to replicate and extend Patil and Duncan’s (2018) demonstration that retrieval establishes an ongoing bias that favors continued associative retrieval. Associative memory accuracy was, indeed, significantly higher following retrieval judgments compared to novelty detection (b = 0.184, SE = 0.064, p < 0.001, 95% CI = [0.059, 0.310]; Fig. 1b); *SI Appendix, Table S1*). This effect held when controlling for the probe trial associate’s stimulus class (i.e. face vs. scene) and the preceding trial’s response time (*SI Appendix, Tables S3, S4 & S4-1*). Further, this effect was not dependent on the associative accuracy of the preceding trial (*SI Appendix, Table S2 & S2-1),* suggesting that this ongoing behavioral bias is more likely initiated by engaging in retrieval rather than the accuracy of the retrieval search. While preceding judgements did not significantly impact response time, participants were marginally slower to retrieve correct associations following retrieval judgments compared to novelty detection (*SI Appendix*, *Table S6*). Replicating Patil & Duncan (2018), preceding memory judgments did not significantly influence item-only recognition memory accuracy (i.e., recognition in the absence of associative retrieval success; t(26) = -1.55, p = 0.134, d = -0.3, 95% CI = [-0.68, 0.09]; Fig. 1c; *SI Appendix, Table S5 & S5-1*). From a theoretical perspective, acts of retrieval have been argued to establish a lingering hippocampal pattern completion bias, facilitating neural reinstatement of other stored information (7, 8, 10, 59). Their selective influence on associative memory accuracy here is consistent with this account. We next sought more direct evidence for this theoretical perspective by asking whether the hippocampus and interconnected regions showed a corresponding bias in memory reinstatement.

### Preceding memory judgments preferentially modulate successive neural reinstatement in high-level ventral stream regions

To test whether retrieval judgments compared to novelty detection biased the hippocampus towards pattern completion, we calculated ERS (14) to measure memory reinstatement. Although hippocampal pattern completion is convincingly demonstrated with invasive recordings (60), it is less consistently detectable with fMRI (61–64). Fortunately, cortical reinstatement—which depends on hippocampal pattern completion (15, 19)—can be robustly measured with fMRI (14, 65). Accordingly, we analyzed probe trial ERS in the hippocampus and high-level ventral stream and frontoparietal ROIs that reliably display memory reinstatement (*a priori:* hippocampus (hipp), medial temporal lobe (MTL) cortex, fusiform cortex (fusiform), medial parietal cortex (mPar); *post-hoc:* dorsal & lateral prefrontal cortex (d&lPFC), ventromedial prefrontal cortex (vmPFC), and lateral parietal cortex (LPC) (64–69)), referred to as *reinstatement ROIs* (Fig. 2a; *SI Appendix, Table S7*). We calculated ERS in three ways, beginning with the most inclusive definition (*broad ERS*). This broad ERS calculation captures the reinstatement of both category information (Scene or Face, necessary for the judgment) and trial-specific information (e.g. Ryan Gosling + tissue). Values above zero indicate reinstatement (see Fig. 2b for schematic and *SI Appendix: Materials & Methods* for calculation specifics).

**Fig. 2.**
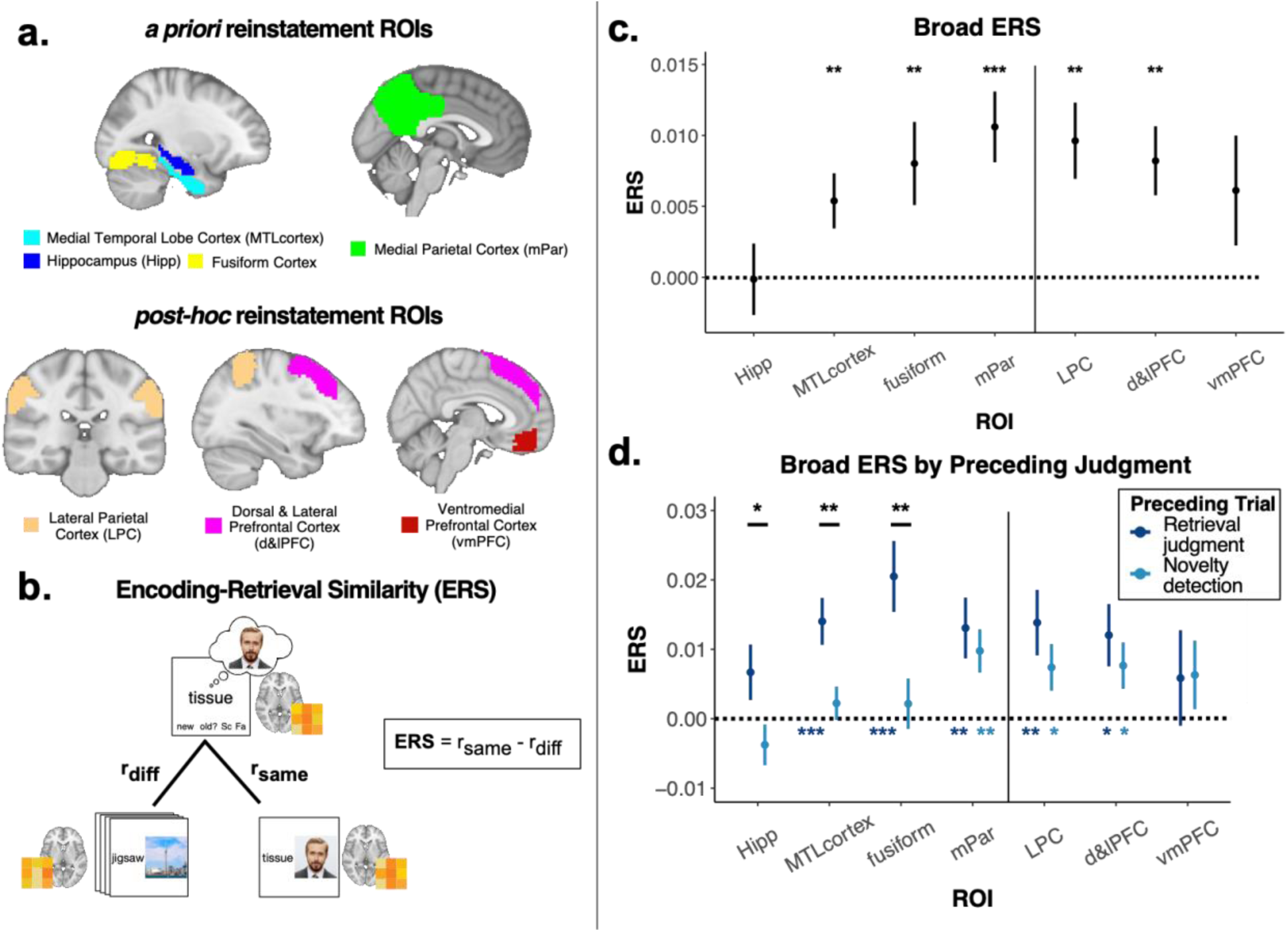
Broad ERS. **a**, *A priori* and *post-hoc* reinstatement ROIs selected for ERS analyses. Probabilistic atlases were used to define structural ROIs (See *SI Appendix, Table S7* for ROI definition details). ***b***, Schematic of broad ERS calculation. Activity patterns from probe trials were correlated with patterns from their corresponding encoding trials (r_same_). r_same_ was corrected by subtracting retrieval trials’ average correlation with encoding trials that included the unprobed image category (r_diff_). This broad ERS calculation captures the reinstatement of both the category of the associate (Face vs. Scene) and the trial-specific information (e.g. Ryan Gosling + Tissue; See *SI Appendix: Materials & Methods*). **c**, Broad ERS on probe trials against chance. Points represent intercepts and error bars represent their standard error from a linear mixed effects model (see *SI Appendix: Materials & Methods*). **d**, Broad ERS on probe trials as a function of preceding memory judgments. Points represent estimated marginal means, and error bars represent their standard error from a linear mixed effects model (see *SI Appendix: Materials & Methods*). Black stars represent significance of the main effect of preceding memory experiences on broad ERS, while dark and light blue stars represent significance of broad ERS against chance following preceding retrieval judgments and novelty detection, respectively. * indicates p < 0.05, ** indicates p < 0.01, *** indicates p < 0.001. Post-hoc ROI p-values are FDR corrected for multiple comparisons.

Medial parietal, MTL, and fusiform cortices all showed significant broad ERS (MTLcortex: b = 0.005, SE = 0.002, p = 0.006, CI = [0.002, 0.009]; fusiform: b = 0.008, SE = 0.003, p = 0.006, 95% CI = [0.002, 0.014]; mPar: b = 0.011, SE = 0.002, p < 0.001, 95% CI = [0.006, 0.015]). The hippocampus did not (b < 0.001, SE = 0.003, p = 0.956, 95% CI = [-0.005, 0.005]; Fig. 2c; *SI Appendix, Table S8*). Importantly, preceding memory judgments influenced ERS in ventral stream regions — notably including the hippocampus — much like they did associative accuracy. Broad ERS was significantly higher following retrieval judgments compared to novelty detection (Hipp: b = 0.005, SE = 0.002, p = 0.027, 95% CI = [0.001, 0.010]; MTLcortex: b = 0.006, SE = 0.002, p = 0.005, 95% CI = [0.002, 0.010]; fusiform: b = 0.009, SE = 0.003, p = 0.003, 95% CI = [0.003, 0.015]; mPar: b = 0.002, SE = 0.003, p = 0.538, 95% CI = [-0.004, 0.007]; Fig. 2d; *SI Appendix, Table S9*). The influence of the preceding memory judgments was so pronounced that broad ERS was only reliably observed in the MTL and fusiform cortices following a preceding retrieval judgment, not novelty detection (following retrieval judgments: estimated marginal mean (EMM_MTL_) = 0.014, p < 0.001, EMM_fusiform_ = 0.021, p < 0.001; following novelty detection EMM_MTL_ = 0.002, p = 0.358, EMM_fusiform_ = 0.002, p = 0.554; p-values FDR corrected for multiple comparisons; Fig. 2d; *SI Appendix, Table S9-1*). These effects held in models accounting for probe trial response time and accuracy (*SI Appendix, Table S10 & S10-1*), suggesting that the reinstatement effects are not secondary to observed behavioural biases. Analyses of frontoparietal *post-hoc* ROIs demonstrated similar trends to those seen in the medial parietal cortex, with the d&lPFC and LPC demonstrating significant broad ERS insensitive to preceding trial judgments (Fig. 2c & 2d; *SI Appendix, Table S8, S9 & S9-1*). In line with our predictions, these results suggest that memory reinstatement is significantly higher across the hippocampus and interconnected high-level ventral stream regions (70) following recent retrieval judgments compared to novelty detection; however, reinstatement in frontoparietal regions is insensitive to these preceding memory judgments.

#### Category & Trial-Specific Reinstatement

Our task also allowed us to distinguish reinstatement of category information (Scene or Face; *category ERS*) from trial-specific information (e.g. Ryan Gosling with tissue*; trial-specific ERS*). Because the task only required scene/face judgments, category-level reinstatement was sufficient for accurate performance. Correspondingly, category ERS was higher for trials with correct compared to incorrect associative memory in the medial parietal cortex (b = 3.856, SE = 1.861, p = 0.038, 95% CI = [0.208, 7.50]; *SI Appendix, Table S11*). Across all ROIs, only the fusiform cortex showed significant category ERS, and the medial parietal cortex demonstrated a marginal effect (fusiform: b = 0.002, SE = 0.001, p = 0.035, 95% CI = [0.000, 0.003]; mPar: b = 0.002, SE = 0.001, p = 0.050, 95% CI = [<0.001, 0.004]; Fig. 3c; *SI Appendix, Table S12*). Fusiform cortex was also sensitive to preceding memory judgments, so much so that it only showed significant category ERS following retrieval judgments (b = 0.004, SE = 0.001, p < 0.001, 95% CI = [0.002, 0.005]; following retrieval judgments: EMM = 0.007, p < 0.001; following novelty detection: EMM = -0.001, p = 0.437; Fig. 3d; *SI Appendix, Tables S13 & S13-1*).

**Fig. 3.**
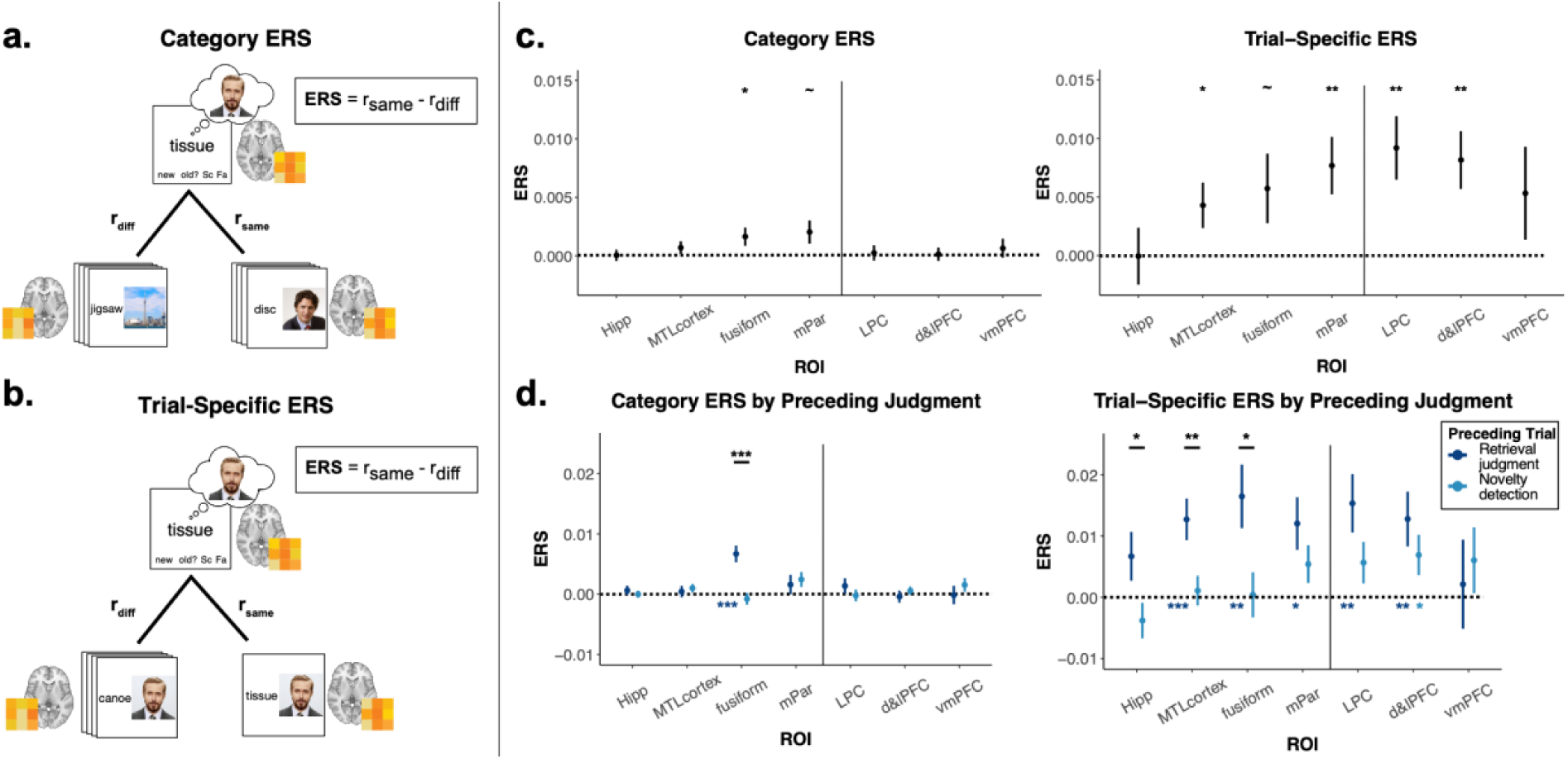
Category and Trial-Specific ERS. **a**, Schematic of Category ERS calculation. Activity patterns from probe trials were correlated with encoding trials that included the unprobed associate from the same image category (r_same_). r_same_ was corrected by subtracting retrieval trials’ average correlation with encoding trials that included the unprobed image category (r_diff_; See *SI Appendix: Materials & Methods*). **b**, Schematic of Trial-Specific ERS calculation. Activity patterns from probe trials were correlated with patterns from corresponding encoding trials (r_same_). r_same_ was corrected by subtracting retrieval trials’ average correlation with all other encoding that included the same associate (r_diff_; *SI Appendix: Materials & Methods*). **c**, Category and Trial-Specific ERS on probe trials against chance. See Fig. 2a and caption for ROI locations, non-abbreviated names, and probabilistic atlas information. Points represent intercepts and error bars represent their standard error from a linear mixed effects model (see *SI Appendix: Materials & Methods*). **d**, Category and Trial-Specific ERS on probe trials as a function of preceding memory judgments. Points represent estimated marginal means, and error bars represent their standard error from a linear mixed effects model (see *SI Appendix: Materials & Methods*). Black stars represent significance of the main effect of preceding memory experiences on category & trial-specific ERS, while dark and light blue stars represent significance of category & trial-specific ERS against chance following preceding retrieval judgments and novelty detection, respectively. ∼ indicates 0.10 > p > 0.05, * indicates p < 0.05, ** indicates p < 0.01, *** indicates p < 0.001. Post-hoc ROI p-values are FDR corrected for multiple comparisons.

Although category ERS on the whole was only modestly modulated by preceding memory judgments, the fusiform cortex’s sensitivity suggests that preceding memory judgments may differentially influence voxels that represent category information. Therefore, we calculated category ERS in participant-specific functional ROIs, consisting of voxels that either responded preferentially to faces or scenes in an independent functional localizer task (see *SI Appendix: Methods & Materials* for full localizer description, ROI definition, and analysis specifications). These functional ROIs reliably reinstated face and scene information (face functional ROIs: b = 0.068, SE = 0.007, p < 0.001, 95% CI = [0.055, 0.081]; scene functional ROIs: b = 0.084, SE = 0.010, p < 0.001, 95% CI = [0.064, 0.103]; *SI Appendix, Table S15*), but this reinstatement was not reliably modulated by preceding memory judgments (face functional ROIs: b = 0.000, SE = 0.001, p = 0.918, 95% CI = [-0.002, 0.002]; scene functional ROIs: b = -0.002, SE = 0.004, p =0.623, 95% CI = [-0.009, 0.006]; *SI Appendix, Tables S16 & S16-1*). Thus, unlike fusiform cortex, structurally unbounded functional ROIs were not sensitive to preceding memory judgments. Combined with the broad ERS findings, these results suggest that sensitivity to preceding memory judgments may depend on a region’s connections with the hippocampus.

We next used trial-specific ERS to identify regions reinstating details beyond stimulus category. Medial parietal and MTL cortices both showed significant trial-specific ERS, while this measure was marginal in fusiform cortex (Hipp: b < 0.000, SE = 0.002, p = 0.984, 95% CI = [-0.005, 0.005]; MTLcortex: b = 0.004, SE = 0.002, p = 0.028, 95% CI = [<0.001, 0.008]; fusiform: b = 0.006, SE = 0.003, p = 0.054, 95% CI = [0.000, 0.012]; mPar: b = 0.008, SE = 0.002, p = 0.002, 95% CI = [0.003, 0.012]; Fig. 3c, *SI Appendix, Table S17*). Preceding memory judgments influenced trial-specific ERS across ventral stream ROIs, notably including the hippocampus (Hipp: b = 0.005, SE = 0.002, p = 0.029, 95% CI = [0.001, 0.010]; MTLcortex: b = 0.006, SE = 0.002, p = 0.005, 95% CI = [0.003, 0.010]; fusiform: b = 0.008, SE = 0.003, p = 0.011, 95% CI = [0.002, 0.014]; mPar: b = 0.003, SE = 0.003, p = 0.209, 95% CI = [-0.002, 0.008]; Fig. 3d, *SI Appendix, Table S18*), with above-chance reinstatement in MTL and fusiform cortices only following retrieval judgments (MTLcortex following retrieval judgments: EMM_MTL_ = 0.013, p = 0.001, EMM_fusiform_ = 0.016, p = 0.004; following novelty detection: EMM_MTL_ = 0.001, p = 0.655, EMM_fusiform_ < 0.001, p = 0.910; p-values FDR corrected for multiple comparisons; Fig. 3d; *SI Appendix, Table S18-1*). As with broad ERS, frontoparietal *post-hoc* ROIs resembled medial parietal cortex, with d&lPFC and LPC showing significant trial-specific ERS insensitive to preceding judgments (Fig. 3d; *SI Appendix, Table S17, S18 & S18-1*). This pattern largely mirrors broad ERS results, suggesting those effects were primarily driven by trial-specific information.

Together, these analyses demonstrate that preceding memory judgments primarily influenced trial-specific reinstatement in hippocampal and high-level ventral stream regions. This pattern is consistent with biased hippocampal pattern completion and associated cortical reinstatement. We next tested whether these reinstatement biases were initiated and mediated through neuromodulatory nuclei responses.

### Neuromodulatory nuclei activity is significantly higher during retrieval judgments compared to novelty detection and predicts subsequent neural reinstatement

We next examined induction trials to identify neural mechanisms that incite reinstatement and retrieval biases. Computational models of hippocampal states predict that novelty detection, compared to retrieval, increases ACh release from the basal forebrain to the hippocampus, biasing processing towards pattern separation (43, 71). This response to novelty detection could reduce subsequent probe trial reinstatement in our paradigm, accounting for the observed reinstatement biases. To evaluate this possibility while remaining open to other mechanisms, we analyzed neuromodulatory nuclei with widespread cortical and hippocampal projections, including the substantia nigra/ventral tegmental area (SN/VTA), locus coeruleus (LC), neocortically projecting basal forebrain (CH4), and hippocampally-projecting basal forebrain (CH123; 44, 63–65). Because induction trials used object stimuli, we also analyzed the anterior hippocampus and perirhinal cortex, regions associated with contextual and object novelty detection (75, 76) (Fig. 4a; *SI Appendix, Table S7)*.

**Fig. 4.**
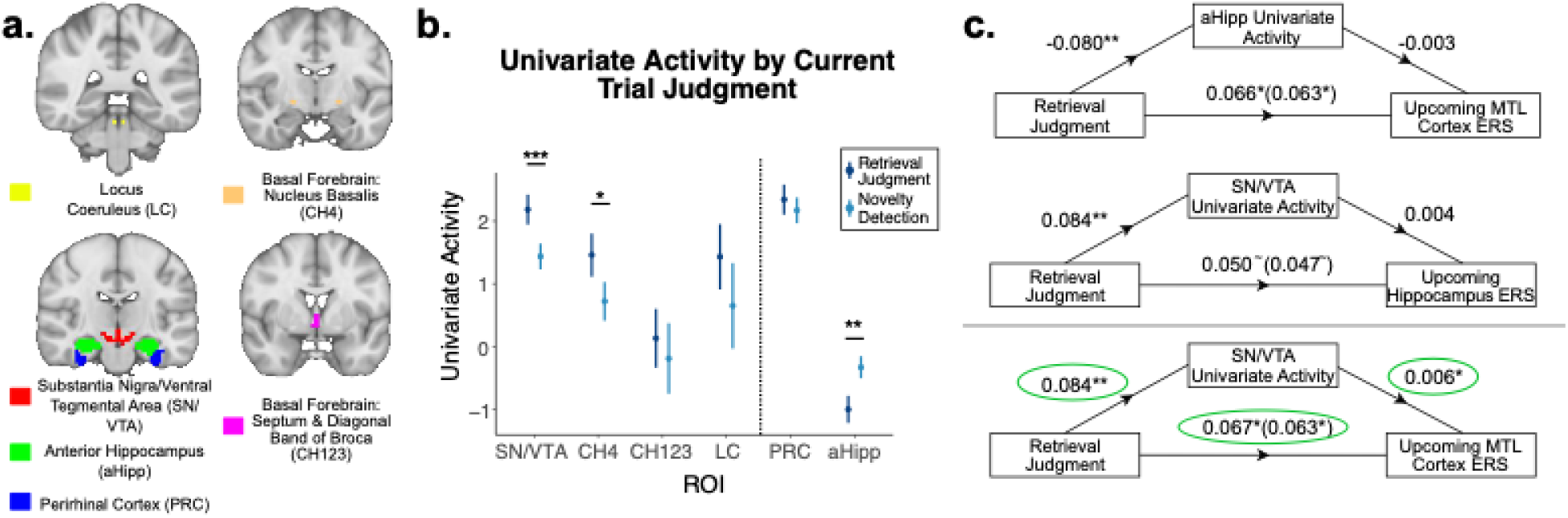
Induction ROI Univariate Activity & ERS Relationships. **a,** Induction ROI labels and locations. Probabilistic atlases were used to define structural ROIs (See *SI Appendix, Table S7* for ROI definition details). **b**, Univariate activity on induction trials separated by memory judgments. Points represent mean beta estimates across participants for retrieval judgment and novelty detection responses. Error bars represent the standard error of the mean, applying the Morey (2008) correction for within-participant comparisons. **c**, Analyses testing whether univariate activity in induction trial ROIs mediates the impact of recent memory experiences (retrieval judgment) on upcoming trial-specific ERS. Models with circled beta values indicate significant partial mediation (See *SI Appendix: Materials & Methods*). ∼ indicates 0.10 > p > 0.05, * indicates p < 0.05, ** indicates p < 0.01, *** indicates p < 0.001.

None of our neuromodulatory ROIs, including the CH123, responded more during novelty detection compared to retrieval judgments. The anterior hippocampus displayed the expected novelty preference (b = -0.092, SE = 0.019, p < 0.001, 95% CI = [-0.130, -0.055]), though it was most active during the implicit baseline. This resulted in negative beta estimates across conditions, which are commonly seen in the hippocampus when passive fixation separates memory trials (77). The perirhinal cortex, LC, and CH123 were insensitive to the manipulation (PRC: b = 0.024, SE = 0.020, p = 0.244, 95% CI = [-0.015, 0.063]; LC: b = 0.020, SE = 0.018, p = 0.232, 95% CI = [-0.013, 0.055]; CH123: b = 0.014, SE = 0.016, p = 0.394, 95% CI = [-0.018, 0.046]).

Most interestingly, the SN/VTA and CH4 both responded more during retrieval judgments compared to novelty detection (SN/VTA: b = 0.085, SE = 0.018, p < 0.001, 95% CI = [0.050, 0.120]; CH4: b = 0.044, SE = 0.018, p = 0.022, 95% CI = [0.009, 0.079]; Fig 4b; *SI Appendix, Table S20*). Given SN/VTA’s and CH4’s established roles in signaling task-relevant goal attainment (78–82), these effects could reflect the motivational or attentional significance of engaging in retrieval compared to detecting novelty in our task. Consistent with this interpretation, attempting to retrieve was sufficient to drive activity in these regions, as neither region’s effect was dependent on the associative accuracy of the retrieval response (*SI Appendix, Tables S21 & S21-1)*. We next explored whether this goal-relevant signaling contributed to upcoming biases in neural reinstatement, and whether it mediated the effect of preceding memory judgments on reinstatement.

We tested if induction trial activity in ROIs sensitive to the task manipulation (CH4, SN/VTA, and anterior hippocampus) predicted successive ERS in the ventral stream ROIs that were biased by preceding memory judgments. We focused on trial-specific ERS because it appeared to drive broad ERS effects (for these analyses applied to broad ERS see *SI Appendix, Table S22*). CH4 induction trial activity did not predict successive ERS, but responses in the two other ROIs did. MTL cortex ERS was weaker following induction trials with high anterior hippocampus activity (b = -0.004, SE = 0.002, p < 0.05, 95% CI [-0.008, 0.000]) and stronger following induction trials with high SN/VTA activity (b = 0.006, SE = 0.002, p = 0.001, 95% CI = [0.002, 0.010]). SN/VTA univariate activity also marginally predicted hippocampal trial-specific ERS (b = 0.005, SE = 0.002, p = 0.058, 95% CI = [0.000, 0.009]; *SI Appendix, Table S23*). In exploratory analyses with reinstatement ROIs that did not demonstrate a preceding memory judgment bias, SN/VTA activity predicted trial-specific ERS in the d&lPFC (b = 0.007, SE = 0.002, p = 0.023, 95% CI = [0.002, 0.012]; p-value FDR corrected across *post-hoc* reinstatement ROIs for multiple comparisons, *SI Appendix, Table S23*). Together, these results suggest that SN/VTA may help prepare ventral and frontal regions to reactivate the details of individual experiences.

Given their induction trial effects and relationship with successive reinstatement, we next asked whether anterior hippocampus and SN/VTA activity mediated the effects of preceding retrieval judgments on trial-specific ERS (see *SI Appendix, Table S24* for broad ERS mediation findings). The mediation analyses did not yield significant findings for two of the three pairs: anterior hippocampus activity did not significantly mediate preceding retrieval judgments’ effect on MTL cortex ERS, nor did SN/VTA activity significantly mediate preceding retrieval judgments’ effect on hippocampal ERS (*SI Appendix, Table S25*). However, SN/VTA univariate activity partially mediated preceding retrieval judgments’ effect on MTL cortex ERS (Fig 4c; *SI Appendix, Table S25*). This finding suggests that preceding SN/VTA activity not only predicts upcoming memory reinstatement but also partially accounts for the power that recent memory judgments exert over memory reinstatement.

To determine whether SN/VTA only influences reinstatement prospectively or also concurrently, we conducted exploratory analyses relating probe trial univariate activity to simultaneous trial-specific ERS. Probe trial activity largely mirrored induction trial effects, with anterior hippocampus showing a novelty preference and CH4 showing a retrieval preference. SN/VTA activity did not significantly differ between probe conditions, though its activity was numerically greater during retrieval judgments (*SI Appendix, Tables S26, S27 & S27-1*). Critically, induction ROI activity did not significantly relate to concurrent broad or trial-specific ERS after correcting for multiple comparisons (*SI Appendix, Table S28 & S29*). However, SN/VTA’s negative relationship with concurrent hippocampal trial-specific ERS was significant before this correction (*SI Appendix, Table S29*), contrasting with its positive predictive relationship with successive ERS. Together, this pattern suggests SN/VTA activity may prepare the brain for future memory reinstatement, rather than facilitating it in the moment.

Finally, given that associative memory accuracy on probe trials, like neural reinstatement, was influenced by preceding memory judgments in our paradigm, we ran a set of analyses to determine whether activity in any of our induction ROIs influenced upcoming behavior in addition to upcoming memory reinstatement. None of our induction ROIs significantly influenced upcoming associative memory accuracy or item-only recognition accuracy (*SI Appendix, Table S30)*. Together, these results suggest that SN/VTA contributes to establishing a bias in neural reinstatement of memories, but not in memory accuracy.

### Activity in dorsal & lateral prefrontal cortex increases during retrieval judgments and predicts successive memory accuracy, but not neural reinstatement

Frontoparietal regions support many cognitive functions involved with ongoing memory retrieval. In addition to their involvement in sustaining attention (83), evidence accumulation (84), and cognitive control (85), several frontoparietal regions have been associated with establishing and sustaining the retrieval mode (55), an ongoing state that allows incoming stimuli to more readily serve as cues for successfully accessing episodic memories (6). Given these functions, we explored whether a set of *frontoparietal induction ROIs*, particularly those involved in establishing the retrieval mode (d&lPFC, frontal pole, mPar, and LPC; 49–56; *SI Appendix, Table S7;* see *SI Appendix: Materials & Methods* for full description of ROI selection), also contributed to establishing biases in neural reinstatement and memory retrieval accuracy. We replicated the full set of analyses applied to our induction ROIs, but here we focus on a subset of findings evaluating the effects of induction trial activity on successive memory reinstatement and behavior (see *SI Appendix, Tables S31-S40* for full set of analyses).

Of the frontoparietal induction ROIs, d&lPFC alone preferentially responded to retrieval judgments on induction trials (b = 0.092, SE = 0.025, p < 0.001, 95% CI = [0.044, 0.142]; *SI Appendix, Table S31*). Therefore, we focus on d&lPFC results here, but report the full set of ROIs in *SI Appendix, Tables S31, S32 & S32-2*. Unlike SN/VTA, d&lPFC activity did not predict successive memory reinstatement (*SI Appendix, Tables S33 & S34*), but did predict successive associative accuracy (b = 0.205, SE = 0.088, p = 0.021, 95% CI = [0.031, 0.378]; *SI Appendix, Table S39*). This d&lPFC induction trial activity did not significantly mediate the impact of preceding memory judgments on upcoming associative accuracy (*SI Appendix, Table S40*). Together, these exploratory findings suggest unique neural contributions to reinstatement and behavioral biases following retrieval judgments.

## Discussion

Here, we explored cognitive and neural mechanisms that contribute to the variability of memory retrieval. First, we replicated prior reports that people are better at retrieving associations following recent retrieval judgments compared to recent novelty detection (13). We found that recent memory judgments biased the fidelity of upcoming memory reinstatement, with high-level ventral stream regions demonstrating significantly higher ERS on trials following retrieval judgments compared to novelty detection. Notably, memory reinstatement in these regions was only reliable following retrieval judgments, not following novelty detection. The SN/VTA arose as the strongest candidate for establishing these neural biases: SN/VTA preferentially responded to retrieval judgments, its activity significantly predicted upcoming reinstatement across ventral stream and frontal ROIs, and its activity partially mediated the effect of recent memory judgments on MTL cortex reinstatement. In a set of exploratory analyses, our d&lPFC ROI, composed of a set of regions associated with attentional control and retrieval mode recruitment, also preferentially responded to retrieval judgments. However, in contrast to the SN/VTA, its activity significantly predicted upcoming associative memory accuracy and not reinstatement. Overall, we uncovered biases in neural reinstatement as a function of preceding memory judgments and a set of underlying neural mechanisms that begin to explain the moment-to-moment variability of memory retrieval.

The manner in which preceding memory judgments shaped reinstatement provides mechanistic insights into retrieval biases. Preceding memory judgments modulated trial-specific reinstatement — the recovery of the individuated experiences — while minimally impacting the coarser category information sufficient for the memory decision. This modulation spanned the high-level ventral stream. The MTL and fusiform cortices both reliably reinstated memories only following retrieval judgments, and while hippocampal ERS did not reliably exceed chance levels, it was significantly stronger following retrieval judgments compared to novelty detection. This modulation of event-specific reinstatement in the hippocampus and highly connected ventral stream regions (70) is consistent with the proposal that recent memory judgements impact associative memory by establishing a pattern completion bias (7–10, 15, 19, 62, 64, 86, 87).

These results also bear on a long-standing puzzle in the literature: fMRI signatures of hippocampal reinstatement are inconsistently observed (61–64), despite this region being the source of associative retrieval (88). Prior work has suggested that this inconsistency is rooted in the hippocampus only reinstating trial-specific details (62, 64, 86, 87), leaving designs that target decontextualized content insensitive to hippocampal episodic representations. Consistent with this proposal, only trial-specific hippocampal ERS was influenced by preceding memory judgments in our task. Our work also points to a second factor: the state dependency of hippocampal ERS. Two of the first studies to find reliable hippocampal reinstatement (89, 90) tested participants’ memory for previously studied associations without interspersing any novelty detection trials; our findings suggest that this continuous focus on retrieval judgments could have amplified hippocampal reinstatement. More broadly, some failures to detect hippocampal reinstatement may reflect the sensitivity of this region to memory states rather than limits on what the hippocampus reinstates, with important implications for future experimental designs.

This study also offered insights into when neuromodulatory nuclei activity may prepare the brain to reactivate memories. Our interest in these nuclei was grounded in demonstrations that their activation can be modulated by motivational factors and salient experiences (24–39), and that the release of neuromodulators can establish lingering effects on memory (33, 36, 39, 41–43, 91). Several studies demonstrated preferential novelty signaling in the SN/VTA, basal forebrain, and locus coeruleus (24–29, 92), leading us to predict stronger responses to novelty detection than retrieval judgments in our task. However, no neuromodulatory nuclei ROIs preferentially signaled novelty detection on induction trials, and the SN/VTA and CH4 preferentially responded to retrieval judgments. This pattern may relate to task demands. SN/VTA activity is broadly linked to goal-maintenance and motivation (78, 79), and CH4 to shifting attentional focus towards motivationally relevant information (80–82). In work demonstrating novelty preferences in these nuclei, participants’ primary task goal was to explore (25–27), detect novel stimuli (24, 28, 29), or was orthogonal to the novelty manipulation (92). In all these cases, trials with novel stimuli could carry motivational salience. By contrast, we used a retrieval task to generate memory states associated with subjective retrieval judgements and novelty detection. Here, participants likely saw their primary goal as retrieving associations, rather than novelty detection, due to both explicit emphasis in instructions and tacit assumptions common to recognition tasks (93, 94). Indeed, studies reporting activity in dopaminergic nuclei and the highly interconnected striatum during retrieval tasks suggest preferential signaling to retrieval judgments, rather than novelty detection (95, 96). Across tasks, then, these nuclei may track the motivational salience of a trial — which happened to coincide with novelty detection in exploration tasks and with retrieval judgments in ours. Future work could more directly test whether the activation of these regions, and its consequences for memory, is governed less by the objective properties of stimuli than by the motivational state they evoke within the context of task goals.

Based on neurocomputational models of the hippocampus (7–9), we predicted that cholinergic, rather than dopaminergic nuclei, would modulate upcoming memory reinstatement. Anchored in rodent physiology, the models propose that increases in hippocampal cholinergic innervation following novelty detection induce a lingering bias away from pattern completion and towards pattern separation (7–9, 36, 97). However, we failed to observe evidence for this mechanism, as the hippocampally projecting CH123 ROI neither preferentially responded to novelty detection nor predicted hippocampal ERS on the following trial. Several methodological limitations, though, caution against an overinterpretation of these null effects. First, CH3—a region that does not strongly project to the hippocampus—was combined with hippocampally projecting CH1 and CH2 regions in the available probabilistic ROI (74). Second, the compositions of these nuclei are heterogeneous, with cholinergic neurons outnumbered by functionally distinct GABA-ergic and glutaminergic neurons (98). Accordingly, while BOLD responses in CH123 should correlate with hippocampal cholinergic innervation, the relationship is indirect and likely weak. By contrast, CH4—the largest basal forebrain nucleus—is composed of approximately 90% cholinergic neurons (99). BOLD activity in this region was sensitive to our task manipulations, showing similar preferences to the SN/VTA, adding to a growing literature that CH4 can be a fruitful target for MRI investigations (100–102). Taken together, while our imaging results did not support the hypothesis that dips in basal forebrain activity prepare the hippocampus to retrieve memories, they also do not undermine decades of relevant research using invasive methods in animals (103–107).

Our imaging results did, however, uncover how the SN/VTA, and potentially resultant dopamine (DA) release, may play a role in preparing the brain to retrieve. Animal research has convincingly demonstrated that DA is critical for the *formation* of memories in the hippocampus; pharmacological and optogenetic activation of hippocampal DA receptors around the time of memory formation enhances later expression of these memories (108–112). By contrast, similar manipulations applied around the time of retrieval have not influenced memory performance (111), leading to the general consensus that DA is important for hippocampal mechanisms underlying memory formation, but not retrieval (113). However, studies that measure or manipulate DA more systemically—rather than targeting the hippocampus—have yielded mixed results. In some human and animal studies, retrieval performance has been positively linked to DA pharmacological manipulations and SN/VTA activity (91, 114–118), suggesting that DA may facilitate retrieval under some circumstances via extra-hippocampal targets. Our finding that SN/VTA predicted reinstatement in the MTL cortex and d&lPFC adds to this work. However, in other studies, no relation was found between DA and retrieval (108, 113, 119–122). Our results suggest a possible reconciliation. SN/VTA responses positively predicted memory reinstatement on successive, but not concurrent, trials. If DA acts to *prepare* upcoming reinstatement rather than to support the retrieval underway, then methods focussed only on current relations may fail to observe how DA influences retrieval. By relating SN/VTA activity to upcoming memory reinstatement, our results point to a potential distinctively preparatory role for DA — one that could be verified with temporally-precise methods for directly measuring and manipulating DA.

We also explored potential preparatory contributions by frontoparietal ROIs previously associated with the retrieval mode. These analyses were exploratory in nature, as our task was not optimized to isolate recruitment of the retrieval mode, defined by greater activity during retrieval attempts (regardless of reinstatement) compared to non-retrieval tasks with similar cognitive demands (55). Our task instructed participants to attempt associative retrieval when they recognized a stimulus and, even when presented with objectively novel stimuli, they may have attempted retrieval (e.g. recall-to-reject, 122). Moreover, we did not have a similarly demanding control task against which to contrast retrieval attempts. Nonetheless, d&lPFC’s preferential response to retrieval judgements over novelty detection may reflect differential recruitment of the retrieval mode across these judgement types. But, because retrieval judgements may also require more attention, evidence accumulation, and cognitive control than novelty detection, the activation pattern could also reflect d&lPFC contributions to these processes more broadly (83–85). Indeed, the frontal pole – a region more specifically tied to the retrieval mode (51) – showed the opposite response, with higher activity during novelty detection than retrieval judgments (*SI Appendix, Table S31*). Regardless, the d&lPFC’s sensitivity to induction trial judgements positioned it to contribute to the mnemonic biases in our task.

Considered together, the SN/VTA and d&lPFC showed a contrasting pattern: SN/VTA activity predicted upcoming reinstatement but not associative accuracy, while d&lPFC predicted accuracy but not reinstatement. This contrast is theoretically suggestive. Tulving distinguished two contributors to successful retrieval: episodic ecphory (reactivation of a stored trace, here indexed through neural reinstatement) and the retrieval mode (a sustained, goal-directed orientation that determines how reinstated information is used) (6). These components need not rise and fall together. One interpretation of our results is that SN/VTA and d&lPFC contribute to the ecphoric and retrieval mode components, respectively. SN/VTA may help prepare brain regions to reinstate detailed, episodic traces. But, ecphory alone may not be sufficient to yield successful retrieval decisions. Aligning with this distinction, trial-specific ERS in the ventral stream did not significantly relate to associative memory accuracy (*SI Appendix, Table S41)*. Instead, associative accuracy was related to concurrent category-level ERS, aligning with the granularity sufficient for the retrieval decision. On the other hand, engagement of the large d&lPFC ROI, consisting of several regions involved in task-relevant attention (83), may have sustained an effective attentional orientation to retrieved content that manifested in an increase in upcoming associative accuracy. We offer this mapping as a theoretical lens, not a fully tested dissociation, to inspire future inquiry. Regardless, our findings uncover a complex set of neural mechanisms that prepare the brain to reinstate the features of memories and support retrieval decisions.

More broadly, this work invites a shift in how we think about the variability of memory retrieval. Retrieval is often treated as a readout of a stored trace — better or worse depending on the quality of the trace or the cue. Our results instead suggest that the brain is continuously shifting in its readiness to retrieve. Each act of remembering may leave the system in a neuromodulatory and representational state that biases how readily the next memory is reactivated, and whether that reactivation is then put to use. The variability of retrieval may therefore reflect, in part, the states we happen to be in when we reach for a memory. We rarely remember in isolation — memories are retrieved in streams, each act of recollection following the last. Our findings suggest that this continuity is not incidental to how well we remember, but a critical determinant of memory success, as the state left by one memory becomes the ground on which the next is retrieved.

## Materials and Methods

Detailed overviews of the methods used in this study are provided in *SI Appendix, Methods and Materials,* including descriptions of our sample, experimental design and procedure, fMRI data processing, and behavioral and fMRI data analysis. Our final analyses included data from 29 individuals convenience sampled from the University of Toronto community. Participants gave written informed consent and were compensated $20-25 CAD/hour for their participation. All experimental procedures were approved by the University of Toronto Research Ethics Board (REB). The experiment consisted of multiple blocks of encoding and retrieval tasks completed within an MRI scanner. Participants studied trial-unique object images paired with one of two repeating words in half of the encoding blocks (object encoding) and studied trial-unique words paired with one of two repeating image categories (face and scene, each of which contained two repeating images) during the other half (word encoding). Stimuli were drawn from publicly available sources (124). During retrieval, participants were shown a trial-unique stimulus, either an object or word, and asked to retrieve the associate shown with it during encoding or flag novel stimuli as new. Retrieval trials were organized in pairs, testing the retrieval of words in response to object images on one trial (induction trials) followed by the retrieval of face/scene images in response to words on the next (probe trials). This retrieval block structure orthogonalized retrieved content across consecutive trials, allowing us to separately analyze object retrieval trials as inductions of memory states and word retrieval trials as probes for measuring the lingering biases evoked from these memory states (see Fig. 1a).

We conducted data analyses in R, using linear mixed effects models to analyze behavioral and fMRI effects of interest. Our main behavioral analyses explored whether associative memory and item-only recognition memory differed as a function of memory judgments made on immediately preceding trials (“retrieval judgment” vs. “novelty detection”). Associative memory accuracy was calculated as the number of correct associative hits out of all associative retrieval attempts (see Fig. 1b), whereas item-only recognition (d’) evaluated participants’ sensitivity to recognize previously seen words when associative recall was not successful (see Fig. 1c). One set of our fMRI analyses mirrored this same question, evaluating whether evidence of neural reinstatement in *a priori* and *post-hoc* ROIs differed as a function of judgments made on preceding trials. We used a set of Encoding Retrieval Similarity (ERS) calculations to estimate neural reinstatement (see Fig. 2b, 3a & 3b for schematics of ERS calculations). We were also interested in whether neural reinstatement or behavioral memory measures may be affected by preceding univariate activity in neuromodulatory nuclei ROIs. We evaluated whether neuromodulatory nuclei ROIs demonstrated significantly higher BOLD responsivity to retrieval judgments vs. novelty detection, then evaluated whether this activity significantly influenced ERS estimates and behavioral memory measures on the immediately subsequent trial. We used mediation analyses to determine whether neuromodulatory nuclei ROI activity significantly mediated the effect that preceding trial judgments had on upcoming ERS and associative accuracy. We additionally explored whether frontoparietal regions played a role in initiating biases in neural reinstatement and behavioral memory measures by replicating the analyses applied to neuromodulatory nuclei ROIs on a set of frontoparietal induction ROIs.

## Resource Availability

All post-processed data and code used to run analyses can be found at https://doi.org/10.17605/OSF.IO/UY9ZV.

## Acknowledgments and Funding Sources

We gratefully acknowledge Justine Vorvis and Hannah Cho for their assistance with data collection, as well as all participants for their time, attention, and willingness to be a part of the research process. This work was supported by the Natural Sciences and Engineering Research Council of Canada (NSERC) Discovery Grant (RGPIN-2023-04231 to Katherine D. Duncan); Canada Research Chair Program (CRC-2023-00363 to Katherine D. Duncan); Ontario Research and Innovation (ER18-14-139 to Katherine D. Duncan); Canada Foundation for Innovation JELF & Ontario Research Fund (34479 to Katherine D. Duncan); and the Ontario Graduate Scholarship (to Anuya Patil).

## Author Contributions

MD led formal analysis, visualization, data curation, writing-draft and writing-review & editing. AP led investigation, methodology, and project administration. AP & MD contributed equally to software. KDD led supervision, funding acquisition, and resources and contributed to writing-editing. AP & KDD led conceptualization. MD, AP, and KDD contributed equally to validation.

