## Supplementary material for "Neural and behavioral catalysts of ongoing memory retrieval": SI Appendix

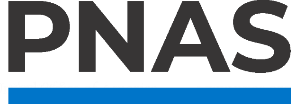

**Supporting Information for**

**Neural and behavioral catalysts of ongoing memory retrieval**

Matthew R. Dougherty^1^, Anuya Patil^1^, Katherine D. Duncan

Corresponding Author: Matthew R. Dougherty

**This PDF file includes:**

Supporting text: Materials & Methods

Tables S1 to S41

SI References

Supporting Information Text

Materials & Methods

***Participants***

Forty-four adults were recruited via convenience sampling from the University of Toronto community through online advertisements and flyers. Two participants withdrew from the study before completing all experiment tasks. Data from 13 participants were excluded from analyses due to previously participating in the study (*n* = 1), showing signs of drowsiness (*n* = 3), failing to follow instructions to stay still inside the scanner (*n* = 1), moving excessively during the study (framewise displacement > 0.3 mm for more than 10% of the volumes on the retrieval and word encoding tasks and 20% of the volumes on the localizer task; *n* = 1), vision issues (*n* = 1), or lower than chance performance on the associative memory task (*n* = 6). Participants were categorized as having lower than chance performance if a binomial test demonstrated that the proportion of correct associative hits out of all associative judgments was not significantly greater than 0.5. The final analyses included 29 participants (21 self-identified females, 7 self-identified males, 1 declined to answer, mean age = 22.9 years [19, 30], mean years of education = 16.43 [13, 22]). Participants identified their race and ethnicity as follows: Japanese/Korean/Chinese (10), Indian/Pakistani/Sri Lankan (8), White (5), Bangladeshi (1), Vietnamese (1), Mixed (Black & White 1, White & Chinese 1), Euroasian (1), and Declined to Answer (1). Participants were right-handed, fluent in English, had normal or corrected-to-normal vision, and had no history of psychiatric, neurological, or learning disorders. Demographic information regarding gender, socioeconomic status, and ancestry were not collected. Convenience sampling resulted in uneven groupings of participants by race/ethnicity and self-identified sex, and as such, we were not sufficiently statistically powered to analyze whether these variables influenced our study’s main results. The racial and ethnic heterogeneity of the University of Toronto community resulted in a diverse sample, increasing generalizability across race and ethnicity. However, the skew of self-identified sex towards female may limit sex-based generalizability. Participants gave written informed consent and were compensated $20-25 CAD/hour for their participation. All experimental procedures were approved by the University of Toronto Research Ethics Board (REB).

***Stimuli***

Stimuli consisted of colored images of 144 common objects (1), two faces of famous individuals (Justin Trudeau, Ryan Gosling, derived from Google Search), two famous Canadian landmarks (CN Tower, Niagara Falls, derived from Google Search), 192 object nouns, and two adjectives (“Plain” and “Safe”). The set of object nouns were selected to not overlap with the images of common objects. For example, if an image of a violin was a part of the 144 object images, the word “violin” was not included in the set of 192 object nouns. Stimuli were presented using PsychoPy (2) on an MR-compatible BOLDscreen monitor (32”, 1920 x 1080) placed behind the scanner. Participants viewed the stimuli through a system of mirrors installed on the head coil. Another image set of objects, faces, places, and scrambled objects were presented in a functional localizer with Matlab & PsychToolbox (see Localizer Tasks).

***Procedure***

Patil & Duncan (2018) demonstrated that the effect of recent memory states on associative memory declines over seconds. Studying this effect necessitates the use of short intertrial intervals (ITIs) during retrieval. However, consistently short ITIs pose challenges for statistically disambiguating responses to consecutive trials due to the slow BOLD response (3). To resolve this issue, we (a) arranged trials into pairs of memory state induction trials and successive probe trials, on which we measured neural reinstatement, (b) fixed the ITIs within trial pairs to 1 second, and (c) orthogonalized content across these two kinds of trials. Specifically, we used images of objects to induce memory states (retrieval and novelty detection). During probe trials, we tested how well participants retrieved a face or a scene when probed with a previously associated word. This alternating pattern, where each pair of trials consisted of independent content, allowed us to estimate how preceding memory states influence successive neural reinstatement (see Fig. 1a). To increase the sensitivity of our reinstatement analyses, we included 5 second fixation periods between retrieval pairs.

The experiment consisted of multiple blocks of encoding, retrieval, and localizer tasks. All tasks were completed inside the MRI scanner. Participants were given instructions and practiced all tasks outside of the scanner until they were comfortable with each task. Participants were aware that their memory for stimuli would be tested before they started the experiment.

***Encoding Blocks***

Participants completed two encoding tasks, corresponding to the induction and probe retrieval trials: object encoding and word encoding, respectively. Their order was counterbalanced across participants.

*Object Encoding*

Participants were shown a series of object images (72 randomly selected from a pool of 144) paired with one of two repeating words, “Plain” or “Safe,” and asked to imagine a scenario in which the word could describe the image. This task was chosen to encourage deep encoding of the object-word pairs. The presentation order of objects was intermixed such that a specific repeating word associate would not repeat across more than three consecutive trials, and the pairing of each object image with “Plain” or “Safe” was randomized for each participant. Each pair was shown for 2.5 seconds. Participants were then given 1.5 seconds to rate the vividness of the association on a 4-point scale of “very vivid” to “not vivid at all,” followed by a 1-second fixation cross. Participants completed one 6.2-minute block of the object encoding task. The memories formed during object encoding were always used to induce memory states during retrieval blocks. fMRI data from the object encoding blocks were not analyzed.

*Word Encoding*

Participants were shown a series of words (96 object-nouns randomly selected from a pool of 192) paired with one of four repeating images of famous landmarks (CN Tower, Niagara Falls) or celebrities (Ryan Gosling, Justin Trudeau). Each image was presented twenty-four times across the two encoding blocks, and the presentation order of images was intermixed such that the image category would not repeat across more than three consecutive trials. The pairing of each word with the repeating images was randomized for each participant. The word was presented on the left side of the screen for 1.5 seconds, followed by one of the four repeating images on the right. The pair together was shown for 2.5 seconds, resulting in 4 second trials. For words paired with a scene, participants were asked to imagine themselves in that scene using that object from a first-person perspective. For words paired with a face, participants were asked to imagine the celebrity using that object against a white background. Participants were then given 1.5 seconds to provide their vividness rating, followed by a 4.5 second fixation cross. Participants completed two 8.2-minute blocks of the word encoding task. Encoding trials in each block were evenly split between scenes and faces. The memories formed during word encoding were used to measure reinstatement during retrieval, and as such, we used a longer ITI than in the object encoding task to facilitate estimates of single-trial fMRI responses.

***Retrieval Session***

Participants completed an associative retrieval session with alternating induction (72 old and 72 new objects) and probe (48 old face-associated, 48 old scene-associated, and 48 new words) trials. Note that we included twice as many old probe trials compared to new because reinstatement could only be measured on old trials. New stimuli were also randomly selected from the same stimulus pools for each participant, excluding the encoded, old stimuli. Trial order was pseudorandomized such that an equal number of “old” and “new” induction trials preceded “old” probe trials, with no more than three consecutive “old” or “new” trials. During each retrieval trial, an object or word was shown on the screen for 2 seconds. Participants were asked to indicate if the object/word was new (by pressing the thumb key) or old, in which case, they were asked to recall the paired associate. For the object trials, the associate responses were Plain (middle finger key) or Safe (ring finger key). For the word trials, the associate responses were Scene (middle finger key) or Face (ring finger key). Participants were asked to press the index finger key if they recognized the object/word but were unable to recall its associate (labelled “old?”). Response options were presented at the bottom of the screen in an intuitive order to help participants remember the mappings. A 1-second and 5-second fixation cross followed each induction and probe trial, respectively. Participants completed four 6.2-minute retrieval blocks.

***Localizer tasks***

After the retrieval blocks, participants completed two functional localizer tasks. We analyzed data from one of these localizers, while the other is beyond the scope of the current project. In the face, place, object (FPO) localizer, participants viewed images of faces, places (synonymous with “scenes” in our paradigm), objects, and scrambled object pictures. There were two runs, each run had four blocks, and there were four mini-blocks in each block (4). Each mini-block was 16 s long, and each image was presented for 500 ms with an ITI of 500 ms. There was a 12-second fixation period between blocks and at the beginning and the end of runs. Participants were asked to press the thumb key when the same image appeared back-to-back. We used data from this localizer task to define functional regions of interest (ROIs) for each participant consisting of voxels that were more responsive to faces than objects and places (face functional ROIs) and voxels that were more responsive to places than faces and objects (scene functional ROIs). We later evaluated patterns of ERS in these functional ROIs.

***Behavioral data analysis***

All analyses were conducted in R (version 4.0.2). Linear mixed effects models were used to reflect the hierarchical nature of our data in which multiple trials are clustered within participants. These models were run with unstructured covariance matrices (lmer function in *lme4* package, version 1.1-23 (5)) with uncorrelated random intercepts and slopes for each within-participant fixed effect varying across participants. We estimated the marginal means of predictors to evaluate their simple effects (*emmeans* package, version 1.11.0 (6)).

We categorized induction trials according to participants’ subjective memory judgments. If they responded with “new,” trials were considered novelty detections; if they recognized the objects, regardless of their associative retrieval success, trials were considered retrieval judgments. We divided probe trials to obtain independent measures of associative memory and item-only recognition memory. To do this, we calculated associative accuracy only on probe trials where participants correctly recognized words as “old,” and item-only recognition memory on trials where participants failed to recall the correct association.

We ran a binomial linear mixed effect model to examine the influence of preceding memory judgments (retrieval judgments vs. novelty detection on induction trials) on successive probe trial associative memory accuracy. Associative accuracy was dummy coded in these models, where correct associative judgments were coded as 1 and incorrect associative and “old?” judgments were coded as 0. For these and all following models, we effect coded the preceding memory judgment predictor (*preceding judgment*: novelty detection as -1; retrieval judgment as 1). We added a covariate to correct for potential response priming, indicating whether the correct associative response on the probe trial was the same as the participants’ response on the preceding trial (*response bias*: dummy coded, unaffected probe trials as 0; affected probe trials as 1). We ran an additional version of this model with preceding judgment as a factor variable with three levels (novelty detection (base level), associative hit (correct associative retrieval judgment), associative miss (retrieval judgment where the incorrect associate was selected)) to explore whether effects significantly differed based on the correctness of the retrieval judgment. We ran post-hoc pairwise comparison t-tests to determine if the levels of the preceding judgment factor variable significantly differed in their effects. In addition, we replicated these models to control for the effects of current trial stimulus class (“Scene” or “Face”, *stimulus class*) and the preceding trial response time (*preceding response time*). Stimulus class was effect coded (“Scene” as -1; “Face” as 1). Preceding response time was z-scored across all induction trials within each participant. We used a paired sample t-test to assess the impact of preceding judgment on item-only recognition accuracy. We calculated *d*′ (z(hit rate) – z(false alarm rate)) for each participant separately for probe trials preceded by novelty detection and retrieval judgments. We ran an ANOVA version of this item-only recognition accuracy analysis with preceding judgment as a factor variable with three levels (novelty detection, associative hit, associative miss) again to explore whether effects differed significantly based on the correctness of the retrieval judgment. We ran post-hoc pairwise comparison t-tests to determine if the levels of the preceding judgment factor variable significantly differed in their effects. Participants were excluded from these analyses of item-only recognition for having hit rates of 0 or false alarm rates of 1, indicating atypically poor performance.

In addition to our behavioral analyses exploring the effect of preceding memory judgments on associative and item-only recognition accuracy, we also ran a linear mixed effect model to examine the influence of preceding memory judgments on accurate probe trial response time. We used an inverse gaussian log linking function to account for the shape of the response time distribution. In addition to preceding judgment, we included four covariates: Response bias, stimulus class, retrieval block number (*centered block number*) and trial number (*scaled trial number;* z-scored within block*)*. The latter two were included to control for task fatigue/learning.

***fMRI data acquisition***

Whole-brain imaging data were acquired on a Siemens Prisma 3T MRI scanner at the Toronto Neuroimaging Facility (ToNI) of the University of Toronto using a 32-channel head coil. Functional images were collected using a T2*-weighted multi-band accelerated EPI sequence (TR = 1.5 s, TE = 26 ms, flip angle = 70º, GRAPPA parallel imaging factor = 2, multiband acceleration factor = 2, distance factor = 20%, field of view = 220 mm, matrix size = 88 x 88, 54 slices oriented parallel to the anterior commissure-posterior commissure line, voxel size = 2.5 x 2.5 x 2.5 mm). One field map was collected (TR = 888 ms, TE 1 = 4.92 ms, TE 2 = 7.38 ms, flip angle = 60º, voxel size = 2.5 x 2.5 x 2.5 mm) to correct EPI images for susceptibility-induced distortions. In addition, one anatomical T1-weighted 3D magnetization-prepared rapid gradient echo (MPRAGE) image (TR = 2 s, TE = 2.67 ms, flip angle = 9º, field of view = 256 mm, matrix = 256 x 256, voxel size = 1 x 1 x 1 mm) was also collected.

***Neuroimaging data processing***

Preprocessing was performed using fMRIPrep 1.2.5 (7, 8), which is based on Nipype 1.1.6 (9, 10).

*Anatomical data preprocessing*

The T1-weighted (T1w) image was corrected for intensity non-uniformity (INU) using N4BiasFieldCorrection (ANTs 2.2.0 (11)) and used as T1w-reference throughout the workflow. The T1w-reference was then skull-stripped using antsBrainExtraction.sh (ANTs 2.2.0), using OASIS as the target template. Spatial normalization to the ICBM 152 Nonlinear Asymmetrical template version 2009c (12, 13) was performed through nonlinear registration with antsRegistration (ANTs 2.2.0 (14)), using brain-extracted versions of both T1w volume and template. Brain tissue segmentation of cerebrospinal fluid (CSF), white-matter (WM) and gray-matter (GM) was performed on the brain-extracted T1w using FAST (FSL 5.0.9 (15, 16)).

*Functional data preprocessing*

The first 4 volumes (6 s) of each of the task BOLD scans were removed to allow for scanner magnet stabilization. For each of the BOLD runs per participant (across all tasks), the following preprocessing was performed. First, a reference volume and its skull-stripped version were generated using a custom methodology of fMRIPrep. A deformation field to correct for susceptibility distortions was estimated based on a field map that was co-registered to the BOLD reference, using a custom workflow of fMRIPrep derived from D. Greve’s epidewarp.fsl script (<https://www.nmr.mgh.harvard.edu/~greve/fbirn/b0/epidewarp.fsl>) and further improvements of HCP Pipelines (17). Based on the estimated susceptibility-induced distortion, an unwarped BOLD reference was calculated for a more accurate co-registration with the anatomical reference. The BOLD reference was then co-registered to the T1w reference using FLIRT (FSL 5.0.9 (18)) with the boundary-based registration (19) cost-function. Co-registration was configured with nine degrees of freedom to account for distortions remaining in the BOLD reference. Head-motion parameters with respect to the BOLD reference (transformation matrices, and six corresponding rotation and translation parameters) were estimated before any spatial transformations using MCFLIRT (FSL 5.0.9 (20)). The BOLD time-series were resampled onto their original, native space by applying a single, composite transform to correct for head-motion and susceptibility distortions.

The BOLD time-series were resampled to MNI152NLin2009cAsym standard space, generating a preprocessed BOLD run in MNI152NLin2009cAsym space. All resamplings were performed with a single interpolation step by composing all the pertinent transformations (i.e. head-motion transform matrices, susceptibility distortion correction, and co-registrations to anatomical and template spaces). Gridded (volumetric) resamplings were performed using *antsApplyTransforms* (ANTs (21)), configured with Lanczos interpolation to minimize the smoothing effects of other kernels (22). Lastly, functional images were further spatially smoothed using a Gaussian kernel of 2mm full-width at half-maximum in FSL.

Several confounding time-series were calculated based on the preprocessed BOLD. Framewise displacement (FD) and DVARS were calculated for each functional run, both using their implementations in Nipype (23). Additionally, a set of physiological regressors were extracted to allow for component-based noise correction (CompCor (24)). Principal components were estimated after high-pass filtering the preprocessed BOLD time-series (using a discrete cosine filter with 128s cut-off) for anatomical CompCor (aCompCor). Firstly, a subcortical mask was obtained by heavily eroding the brain mask, which ensured it did not include cortical grey matter (GM) regions. Six aCompCor components were then calculated within the intersection of the aforementioned mask and the union of cerebrospinal fluid (CSF) and white matter (WM) masks calculated in T1w space, after their projection to the native space of each functional run (using the inverse BOLD-to-T1w transformation). The head-motion estimates calculated in the correction step were also placed within the corresponding confounds file.

Many internal operations of fMRIPrep use Nilearn 0.5.0 (25), mostly within the functional processing workflow. For more details of the pipeline, see the section corresponding to workflows in fMRIPrep’s documentation (<https://fmriprep.org/en/1.2.5/workflows.html>).

***ROI selection***

To measure reinstatement, we defined four *a priori* ROIs (Fig. 2a) that have previously been shown to reinstate trial-specific and category information (26–29): whole hippocampus, medial temporal lobe cortex (MTLcortex), fusiform cortex (fusiform), and medial parietal cortex (mPar). After conversations with colleagues, we added three additional frontoparietal *post-hoc* ROIs (Fig. 2a) that have previously been shown to reinstate trial-specific or category information: lateral parietal cortex (LPC), dorsal & lateral prefrontal cortex (d&lPFC), and ventromedial prefrontal cortex (vmPFC) (26–31). We refer to all seven of these regions as *reinstatement* *ROIs*. We additionally defined and analyzed reinstatement in a set of participant-specific functional ROIs based on our FPO localizer. To do this, we ran general linear models (GLMs) to localize BOLD signal specific to different stimulus contrasts during the FPO localizer task: preferential signaling of faces over scenes and objects (faces effect coded to +1, scenes and objects each effect coded to -0.5) and preferential signaling of scenes over faces and objects (scenes effect coded to +1, faces and objects each effect coded to -0.5). We defined functional ROIs of equivalent size across participants consisting of the most sensitive voxels for each contrast. We restricted the included voxels to those in the whole brain mask, however, anatomical location of these voxels within the brain was not relevant to the line of questions guiding this analysis. As such, we did not otherwise anatomically constrain these functional ROIs. Following previous research (32–34), we determined the most sensitive voxels through a top percentage cutoff of the beta estimates from the GLMs (top 0.25%) that made these functional ROIs a similar volume (1179 voxels) to the median of our *a priori* and *post-hoc* reinstatement ROIs (fusiform ROI; 1367 voxels).

To measure responsivity to novelty detection and retrieval judgments during the induction trials, and potential relationships to upcoming probe trial reinstatement and behavior, we evaluated six *a priori induction ROIs* (Fig. 4a): basal forebrain medial septum and the diagonal band of Broca (CH123), basal forebrain nucleus basalis (CH4), substantia nigra and ventral tegmental area (SN/VTA), locus coeruleus (LC), anterior hippocampus (aHipp), and perirhinal cortex (PRC). Upon analyzing and interpreting data from our induction ROIs, we elected to define a set of *post-hoc frontoparietal induction ROIs* to understand whether and how they may also contribute to establishing retrieval processing biases in memory reinstatement and behavior. We selected four ROIs based on previous empirical work showing activity in these regions tracks retrieval mode recruitment (35–42). Many of these regions overlapped with our reinstatement ROIs, and as such, we utilized the same masks where applicable (medial parietal cortex (mPar), lateral parietal cortex (LPC), dorsal & lateral prefrontal cortex (d&lPFC)) and defined the final region (frontal pole (FP)) using the same probabilistic atlas for consistency (Harvard-Oxford Probabilistic Atlas). The probabilistic atlases and thresholds used for each brain region are listed in SI Appendix, Table S7. All masks were defined in T1 MNI space and resampled to the functional MNI space.

***Generating Single Trial Estimates***

GLMs were constructed for each of the 96 word encoding and 288 retrieval trials to estimate single-trial beta values. This was done using the LS-S method (43) implemented in *FSL* with *Nipype* (version 1.8.5) and in custom Python (version 2.7) scripts in which separate GLMs were conducted for each trial. These GLMs included a regressor for the trial of interest, and responses to all other trials were modelled together with a second regressor. Each trial was modelled at stimulus onset (for word encoding, the time when the word and the image were first displayed together) using a 2.0-second (retrieval task) or a 2.5-second (word encoding task) boxcar function convolved with a double-gamma hemodynamic response function. In addition to these primary regressors of interest, each GLM also included the following nuisance regressors: the temporal derivatives of six motion estimates and six aCompCor motion components. Volumes for which the framewise displacement was greater than 0.3 mm were added as spike regressors in the GLMs.

***fMRI data analysis***

We obtained three estimates of neural reinstatement that differed in their levels of specificity. The first (broad ERS) estimated both category information about the associate (required to make a retrieval judgment; “Scene” vs. “Face”), as well as specific information about the trial itself (the specific scene/face and its association with the probed word). The second (category ERS) estimated only the reinstatement of the associate category, while the third (trial-specific ERS) estimated only the reinstatement of each trial’s specific association. For each ERS measure, we calculated two Pearson correlations and operationally defined ERS as the difference between them (r_same_ – r_diff_). For broad ERS, activity patterns from probe trials were correlated with patterns from corresponding encoding trials (r_same_) and corrected by subtracting retrieval trials’ average correlation with encoding trials that included the unprobed image category (e.g., scenes if the retrieval trial probed a face; r_diff_; Fig. 2b). For category ERS, activity patterns from probe trials were correlated with encoding trials that included the unprobed associate from the same image category (e.g. if the retrieval probe was studied with Ryan Gosling, all Justin Trudeau encoding trials; r_same_) and corrected by subtracting retrieval trials’ average correlation with encoding trials that included the unprobed image category (e.g., scenes if the retrieval trial probed a face; r_diff_; Fig. 3a). For trial-specific ERS, activity patterns from probe trials were correlated with patterns from corresponding encoding trials (r_same_) and corrected by subtracting retrieval trials’ average correlation with all other encoding that included the same associate (e.g., if tissue were the probe, all Ryan Gosling encoding trials except for Ryan Gosling + tissue; r_diff_; Fig. 3b). For all ERS analyses, we Fisher z-transformed correlation coefficients and only included probe trials on which participants correctly recognized words as “old.”

We ran a linear mixed effect model for each ERS measure to examine whether regions significantly reinstated learned information. We included response bias as a covariate in all ERS analyses, as it significantly related to ERS in several reinstatement ROIs (SI Appendix, Table S8).

We tested whether preceding judgments (novelty detection vs. retrieval judgments) influenced ERS on probe trials in each of the reinstatement ROIs using linear mixed effects models. Preceding judgment was the primary predictor and response bias was included as a covariate. We ran an additional model for each ROI to control for the effects of the *accuracy* and *response time* on the current trial. Accuracy was effect coded (accurate trials as 1; inaccurate trials as -1), while response time was z-scored across all probe trials for each participant.

We were interested in whether category ERS significantly related to participants’ associative accuracy, as category information was needed for correct judgments (“Scene” vs. “Face”). We ran a generalized linear mixed effects model for each of our *a priori* reinstatement ROIs predicting whether associative accuracy significantly related to category ERS. Each model included Fisher z transformed category ERS as the primary predictor as well as response bias and preceding judgment as covariates. We replicated these models in an exploratory analysis evaluating whether trial-specific ERS significantly related to associative accuracy.

To evaluate whether induction ROIs preferentially responded to novelty detection or retrieval judgments, we ran linear mixed effects models predicting average univariate signal, z-scored within participant. Each model included the current trial judgment (*judgment*) as a predictor, effect coded with novelty detection as -1 and retrieval judgments as 1. We ran additional versions of these models with judgment as a factor variable with three levels (novelty detection (base level), associative hit, associative miss) to explore whether effects differed significantly based on the correctness of the retrieval judgment. We ran post-hoc pairwise comparison t-tests to determine if the levels of the judgment factor variable significantly differed in their effects. We ran additional follow up analyses exploring this on probe trials using models with identical structure. We applied this same set of analyses to our *post-hoc* frontoparietal induction ROIs.

To determine whether induction trial univariate signal in our induction ROIs predicted successive reinstatement, we ran linear mixed effect models predicting trial-by-trial ERS estimates in each of our reinstatement ROIs. Each model included univariate signal on the preceding induction trial in each of the induction ROIs, z-scored within participant, and response bias as predictors. We ran additional follow up analyses using models with identical structure evaluating the relationship between concurrent probe trial univariate activity in induction ROIs and ERS, rather than successive relationships. We applied this same set of analyses to our *post-hoc* frontoparietal induction ROIs, but only analyzed their relationship to a subset of reinstatement ROIs whose ERS was significantly modulated by preceding judgments.

We then performed multilevel mediation analyses using *lme4* (version 1.1-23) and *mediation* (version 4.5.0) packages in R (version 4.0.2) to assess if univariate signal in induction ROIs (mediator) mediated the effects of preceding judgments (independent variable; IV) on successive ERS (outcome). To establish mediation, there must be evidence for all pair-wise relations. Only three pairs of ROIs met these criteria: SN/VTA induction trial activity as a mediator for preceding judgment on hippocampal and MTL cortex ERS, as well as anterior hippocampus induction trial activity as a mediator for preceding judgment on MTL cortex ERS (ultimately, these models were run for both broad and trial-specific ERS estimates). The following multilevel models were run as part of the mediation:

Model (1): Influence of preceding judgment on univariate activity in induction ROI (mediator ~ IV)

Model (2): Influence of preceding judgment and univariate activity in all of the induction ROIs on successive ERS (outcome ~ IV + mediator).

These models were run on data that included correctly recognized probe trials only. Response bias was included as a covariate in Model (2). Both univariate activity and ERS estimates were z-scored within participant for these models.

Finally, we performed a multilevel mediation analysis to assess if univariate signaling in frontoparietal induction ROIs (mediator) mediated the effects of preceding judgments (IV) on successive associative accuracy (outcome). Only two ROIs demonstrated significance for all pair-wise relations: d&lPFC and mPar. The following multilevel models were run as part of the mediation:

Model (1): Influence of preceding judgment on univariate activity in frontoparietal induction ROI (mediator ~ IV)

Model (2): Influence of preceding judgment and univariate activity in all of the frontoparietal induction ROIs on successive associative accuracy (outcome ~ IV + mediator).

These models were run on data that included correctly recognized probe trials only. Response bias was included as a covariate in Model (2). Both univariate activity and ERS estimates were z-scored within participant for these models.
